# Enhancing the prediction of protein coding regions in biological sequence via a deep learning framework with hybrid encoding^★,★★^

**DOI:** 10.1101/2020.11.07.372524

**Authors:** Chao Wei, Junying Zhang, Xiguo Yuan

## Abstract

Protein coding regions prediction is a very important but overlooked subtask for tasks such as prediction of complete gene structure, coding/noncoding RNA. Many machine learning methods have been proposed for this problem, they first encode a biological sequence into numerical values and then feed them into a classifier for final prediction. However, encoding schemes directly influence the classifier’s capability to capture coding features and how to choose a proper encoding scheme remains uncertain. Recently, we proposed a protein coding region prediction method in transcript sequences based on a bidirectional recurrent neural network with non-overlapping 3-mer feature, and achieved considerable improvement over existing methods, but there is still much room to improve the performance. First, 3-mer feature that counts the occurrence frequency of trinucleotides in a biological sequence only reflect local sequence order information between the most contiguous nucleotides, which loses almost all the global sequence order information. Second, kmer features of length *k* larger than three (e.g., hexamer) may also contain useful information. Based on the two points, we here present a deep learning framework with hybrid encoding for protein coding regions prediction in biological sequences, which effectively exploit global sequence order information, non-overlapping gapped kmer (gkm) features and statistical dependencies among coding labels. 3-fold cross-validation tests on human and mouse biological sequences demonstrate that our proposed method significantly outperforms existing state-of-the-art methods.

## 1. Introduction

Genome annotation helps in understanding complicated biological mechanisms underlying gene regulation and remains a challenging problem in biology. The development of next-generation sequencing (NGS) technologies give rise to an exponential increase of sequence data. Many efforts have been dedicated to the identification of genomic mutations by using NGS datasets Yuan, Zhang, Yang, Bai and Fan (2017); Tuo, Liu and Chen (2020) in the past few years, it is urgent to find effective genome annotation techniques for predicting genes Catherine, Marie-France, Thomas and Pierre (2002).

The prediction of protein coding regions in genomic or transcript sequences is a very important but overlooked subtask for genome annotation. Many well-known gene prediction tools (e.g., GenScan Burge and Karlin (1997), Augustus Stanke, Steinkamp, Waack and Morgenstern (2004)) are integrated models, in which the task of identifying gene structure is first divided into subtasks such as the prediction of functional sites and coding regions, and then these subtasks are integrated into a structured learning framework for the prediction of gene structure Al-Turaiki, Mathkour, Touir and Hammami (2011). Moreover, coding features is also very important for computational methods to discriminate mRNAs from long non-coding RNAs Li, Zhang and Zhou (2014); Tong and Liu (2019). However, prediction of protein coding regions from uncharacterized biological sequences (e.g., genomic or transcript sequences) is a very challenging task. This is because (1) genomic sequences contain introns that disrupt the coding structure Catherine et al. (2002). (2) there exists a considerable number of short exons bordered by large introns, which is easily missed by computational methods Catherine et al. (2002). (3) unlike consensus motifs, coding features often exhibit higher-order distant interactions among nucleotides and more difficult to capture Rajapakse and Ho (2005).

Many existing computational methods Hatzigeorgiou, Mache and Reczko (1996); Guigó (1997); Zhang, Lin, Yan and Zhang (1998); Hatzigeorgiou (2002); Shuo and Yi-sheng (2009); Tzanis, Berberidis and Vlahavas (2012); Wei, Zhang, Yuan, He, Liu and Wu (2020) have been proposed for protein coding regions prediction in genomic or transcript sequence during the past decades. They first encode a biological sequence into numerical values and then feed them into a classifier for final prediction. There are mainly two types according to the encoding scheme they use: sequential model and discrete model. The sequential model converts each nucleotide of a biological sequence into a numerical value one by one, which preserves the original order of the bases that appears in the biological sequence. A widely used encoding scheme is one-hot representation (also called C4 encoding) Voss (1992) that encodes four nucleotides with a binary vector of four bits (A-[1,0,0,0], C-[0,1,0,0], etc.). The binary numbers for each nucleotide are orthonormal to each other and have identical Hamming distance. The one-hot encoding scheme is not only applied to protein coding regions prediction, but also a large number of applications Alipanahi, Delong, Weirauch and Frey (2015); Min, Zeng, Chen, Chen, Chen and Jiang (2017); Du, Yao, Diao, Zhu, Zhang and Li (2018); Zuallaert, Kim, Soete, Saeys and Neve (2018). In contrast, the discrete model exerts efforts on engineering a set of features based on prior knowledge from a biological sequence. Some widely used biological features include the codon usage Staden and McLachian (1982), codon prototype Shepherd and J. (1981), hexmer usage Claverie, Sauvaget and Bougueleret (1990), and Z curves of biological sequence Chun-Ting and Ren (1991), which have been comprehensively reviewed by Fickett and Tung (1992).

The abovementioned two models have both merits and demerits. In fact, data representation in genome analysis plays an important role in computational methods to learn relevant biological features Yu, Li and Yu (2018); Kalkatawi, Magana-Mora, Jankovic and Bajic (2019), however, effectively encoding biological sequences and building computational methods for features learning remains uncertain Yu et al. (2018). The sequential model preserves the global sequences order information Chen, Feng, Deng, Lin and Chou (2014) but computational methods could not fully capture biological features by this model. As mentioned by Rajapakse and Ho (2005); Li, Liu, Wong and Yap (2004), it is not easy for neural networks to learn high-order correlations from extremely low-level inputs, e.g., a string of nucleotides. The work Fu, Peng and Chai (2019) also claim that sequential model like one-hot encoding may contain limited useful information compared to other objects like images or sounds that is more suitable for deep learning. Recently, Choong and Lee (2017) claims that one-hot encoding is unable to capture the frequency domain of features like kmer. On the contrary, a discrete model like 3-mer representation of biological sequences is a feature that has proved to be a successful means to discriminate between coding and non-coding regions for the fact that the distribution over the 64 different codons is significantly different in coding regions compared to non-coding regions Axelson-Fisk (2010). Despite the effectiveness of 3-mer, it can only incorporate local sequence order information between the most contiguous nucleotides and none of the global sequence order information can be reflected Chen et al. (2014). Moreover, kmer features of different length *k* (e.g., hexamer) may also be useful for coding potential prediction Guigó (1997); Li et al. (2014).

Based on the aforementioned analysis, we explore how to enhance the prediction of protein coding regions in genomic and transcript sequences by integrating sequential with the discrete model. We propose a novel method for protein coding regions prediction by using a hybrid convolutional neural network Lecun, Bengio and Hinton (2015) and bidirectional recurrent neural network Schuster and Paliwal (1997) framework (CNN-BRNN), which effectively exploits global sequence order information, non-overlapping gapped kmer (gkm) features, and statistical dependencies among coding labels. Evaluated on genomic and transcript sequences, our method gives an excellent prediction performance, which significantly outperforms existing state-of-the-art method. There are three contributions which may explain the excellent performance of our proposed framework:

- We present a CNN-BRNN framework for protein coding regions prediction both in genomic and transcript sequences, which significantly outperforms existing state-of-the-art methods.
- We exploit a hybrid encoding (e.g., C2 Arniker, Kwan, Law and Lun (2011) and gkm Ghandi, Lee, Mohammad-Noori and Beer (2014)) for protein coding regions prediction for the first time, it fuses global sequence order information and kmer features simultaneously, which demonstrate improved prediction performance over using each single encoding.
- Inspired by our previous work Wei et al. (2020), we extend label dependencies to genomic sequences and significantly improve the prediction performance on genomic sequences over existing methods.

The source code and the dataset used in the paper are publicly available at: https://github.com/xdcwei/DeepCoding/.

## 2. Related works

We here review the most relevant works to us.

### Hybrid encoding

A few previous works demonstrate that combining sequence information with biological features can bring considerable performance improvement in specific applications. It is firstly introduced for promoter prediction in Xie, Wu, Lam and Yan (2006), who proposes a method called PromoterExplorer which integrates various biological features (e.g., local distribution of pentamers, positional CpG island features) with digitized DNA sequence in a cascade AdaBoost-based classifier and achieves promising performance. The work Chen et al. (2014) presents a sequence-based predictor, called iTIS-PseTNC, for identifying translation initiation site in human genes and claims that using kmer representation of DNA sequences only reflect local sequence order information but lose all global sequence order information. They Chen et al. (2014) remedy this by using a collaborative representation called pseudo trinucleotide composition which incorporates the physicochemical properties into DNA sequence and combines with kmer features. Recently, in the work of Fu et al. (2019), they propose a hybrid sequence-based deep learning model called MHCpG, which integrates MeDIP-seq data with Histone information to predict DNA methylated CpG states, it exceeds the other approaches and gained more satisfactory promoter prediction performance owing to fusing multiple biological relevant features and sequence information.

### Gapped kmer

kmer is a simple but effective feature that has been successfully applied to bioinformatics, e.g., the prediction of protein coding regions Staden and McLachian (1982); Hatzigeorgiou (2002), coding potential Li et al. (2014); Tong and Liu (2019), regulation elements Ghandi et al. (2014); Fu et al. (2019). However, it suffers from the inherent limitation that the increase of *k* leads to a very long and sparse feature vector Ghandi et al. (2014). To address this problem, Ghandi et al. (2014) introduces a concept of gaps that is defined as mismatches exist in kmer. Gapped kmer (gkm) is not only effective at reducing the length and sparseness of feature vector, but also biological significant–as the results of changes during the evolution process, initial and resultant biological sequences still share many common features in spite of sequence dissimilarities Wang, Xu and Liu (2016b). There are a considerable number of works demonstrate that gkm can achieve improved prediction performance over kmer Ghandi et al. (2014); Wang et al. (2016b).

All the above works give us a strong intuition that we can enhance the prediction of protein coding regions by incorporating global sequence order information and biological features like gkm Ghandi et al. (2014).

## 3. Materials and Method

In this section, datasets, definitions of problems, data representation of biological sequences, the CNN-BRNN framework for protein coding regions prediction are introduced. The graphical illustration of the proposed method is shown in Figure 1.

**Figure 1:**
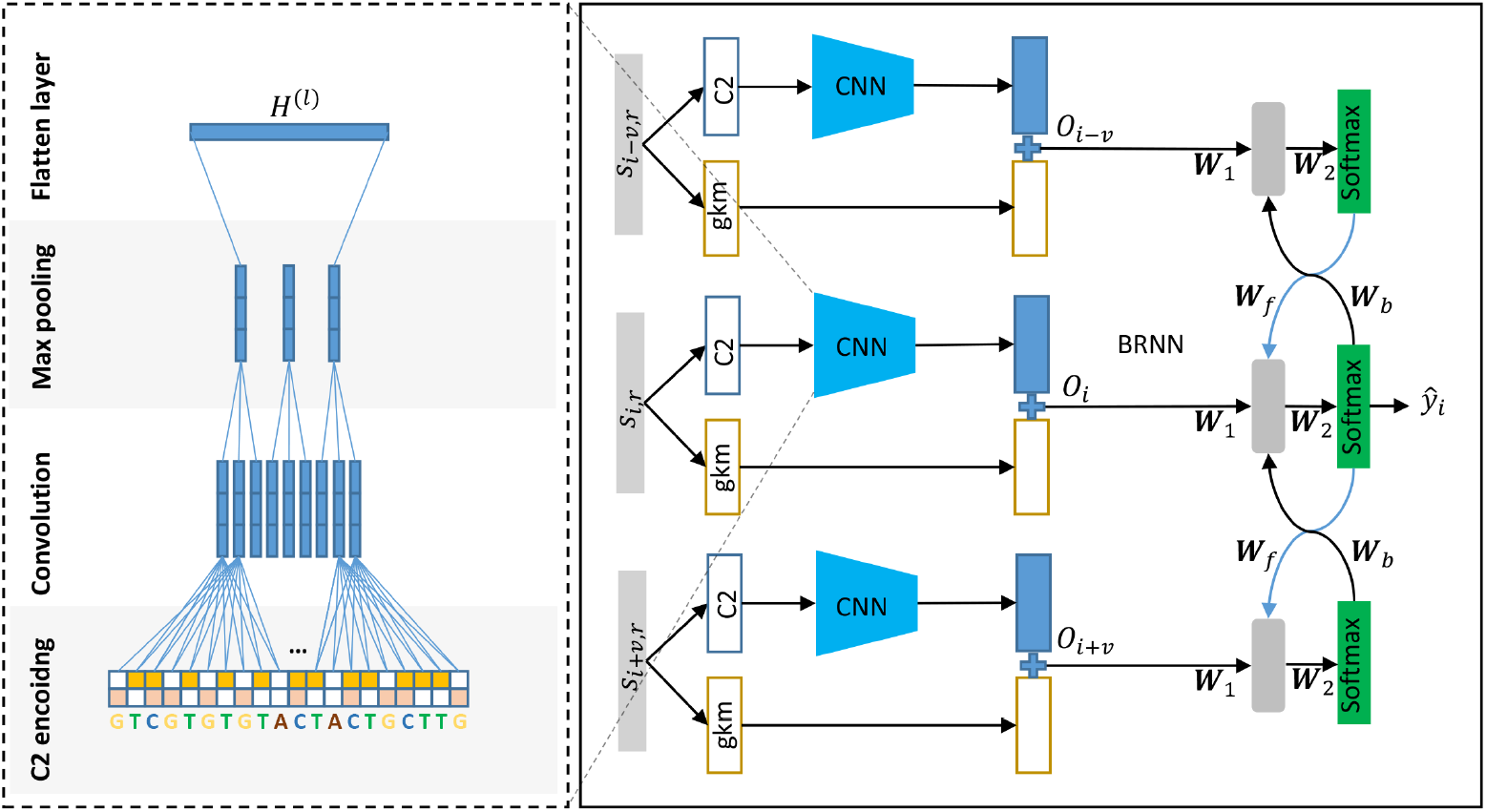
A graphical illustration of the proposed CNN-BRNN architecture for protein coding regions prediction. For each position in a biological sequence, the current subsequence and its neighboring subsequences are firstly encoded into C2 and gkm encoding, then C2 encoding into a CNN and merges with gkm, which are finally fed into a BRNN for protein coding regions prediction.

### 3.1. Datasets

As shown in Table 4, we bulid genomic and transcript datasets of human and mouse from Refseq Pruitt, Tatusova and Maglott (2007) that provides comprehensive, non-redundant and well-annotated set of sequences. For genomic datasets, only one isoform is randomly selected from alternative isoforms of the same gene. A total number of 19,288 and 16,473 sequences are obtained for human and mouse dataset, respectively. For transcript datasets, mRNA sequences with prefixes ‘NM_’ is selected. A total number of 24,842 and 19,900 sequences are obtained for human and mouse dataset, respectively. Then coding samples are selected from all the biological sequences and randomly shuffled. To avoid the imbalanced data problem, negative examples are chosen such that their number equals that of the positive examples. Finally, all the samples are split into 3 parts to perform 3-fold cross-validation and similar samples are removed from test data to guarantee that each sample in test data has no more than 50% identity with any sample in training data.

### 3.2. Preliminaries

In what follows, *s* = *s*_1_*s*_2_…*s*_*n*_ is a biological sequence (e.g., DNA or mRNA), where *s*_*i*_ ∊ {*A, C, T, G*}, and *y* = *y*_1_*y*_2_…*y*_*n*_ is the label sequence of *s*, where *y*_*i*_ ∊ {1, 0} denote the position *i* in *s* is coding (*y*_*i*_ = 1) or not (*y*_*i*_ = 0). Then the protein coding regions prediction is equivalent to solve the following *maximum a posteriori* (MAP) estimation problem

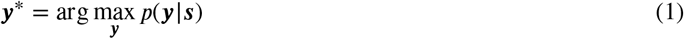

Almost all previous machine learning methods regard protein coding regions prediction as an independent binary classification problem and adopt a sliding window strategy to discriminate coding or non-coding, then the Eq. 1 can be formulated as:

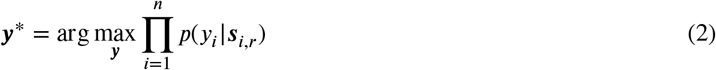

and the predictions can be made separately, in the form

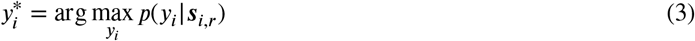

where *s*_*p,r*_ indicate a subsequence of *s* centered at position *p* with a fixed length window 2 × *r* + 1. In our previous work, we demonstrate the significance of exploiting label dependencies among coding labels and improve the prediction performance in transcripts. In this work, we also extend the label dependencies to genomic sequences. Hence, for a position *i* in a genomic or transcript sequence, we here consider the following MAP problem:

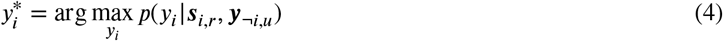

where ¬ denote not operator and *y*_¬*i,u*_ = *y*_*i−u*_…*y*_*i*−1_*y*_*i*+1_…*y*_*i*+*u*_. It can be observed from Eq. 4 that whether the position *i* in ***s*** is coding or not depends on not only its local region ***s***_*i,r*_, but also its neighboring coding labels ***y***_¬*i,u*_. This characteristic resembles the linear-chain conditional random fields (CRF) Lafferty, Mccallum and Pereira 2001) that encodes state features and transition features. State features encode the content properties of the current position, while transition features focus on state transition information (e.g., in our work, coding to coding or non-coding to non-coding). It is worth emphasizing that the main difference of our model from CRF is that, we here consider long-range dependencies between labels while CRF consider the most neighboring two labels (e.g., *y*_*i*_ and *y*_*i*−1_). In the following subsection, we introduce a CNN-BRNN architecture to effectively estimate conditional probability *p*(*y*_*i*_|***s***_*i,r*_, *y*_¬*i,u*_).

### 3.3. CNN-BRNN for protein coding regions prediction

#### 3.3.1 Hybrid encoding

The basic problem of machine learning methods for protein coding regions prediction is designing an effective encoding scheme for a biological sequence. Considering the demerits of the sequential model and discrete model, we propose a hybrid encoding scheme that combines the sequential model and discrete model, which could exploit the joint merits of each model. Given a subsequence ***s***_*i,r*_, the hybrid encoding can be represented as:

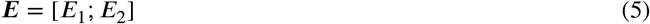

we here adopt a sequential model such as C2 Arniker et al. (2011) to capture global sequence order information and hence *E*_1_ can be formulated as:

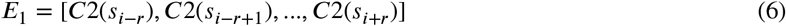

where *C*2 converts a nucleotide into 2-bit binary (e.g., A-[0,0], C-[1,1],G-[1,0],T-[0,1]), note that C4 encoding Arniker et al. (2011) is also a sequential model that preserves the global sequence order information, however, it is more computationally expensive than C2 encoding, hence we here adopt C2 encoding to substitute for C4 encoding. Meanwhile, the sliding window size is relevant to prediction performance, the use of a smaller size can increase the accuracy on the border of exon but leads to of higher rate of false positive prediction Hatzigeorgiou et al. (1996). They Hatzigeorgiou et al. (1996); Shuo and Yi-sheng (2009) use 91 whereas we adopt 90 in practice for convenience of counting the number of codons. The slight difference has almost no effect on the prediction performance.

As for discrete model, we adopt a non-overlapping gkm Ghandi et al. (2014) to capture local sequence order information. There are two parameters for gkm. (1) the whole word length *l* and (2) the number of non-gapped (informative) positions *k*. The number of gaps is thus *l* − *k*. We here set *l* = 5 and *k* = 3, which not only effectively reduce the dimensions of feature vector from 4^5^ = 1024 to 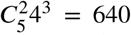, but also contain non-overlapping 3-mer information (e.g., *AAAXX, …, T T T XX*). Hence, *E*_2_ can be formulated as:

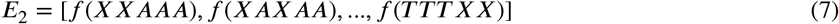

where *f*(*XXAAA*) counts the occurrence frequency of non-overlapping gapped trinucleotides *XXAAA* in biological sequence. By introducing two gaps *XX*, two words *GT GCA* and *CT ACA* of length 5 have the same gapped trinucleotides *XT XCA*.

#### 3.3.2. CNN-BRNN

Given hybrid encoding for each sliding window ***s***_*i,r*_, another problem is how to build an effective machine learning method to integrate global sequence order information, non-overlapping kmer features, and label dependencies. We present a hybrid CNN-BRNN architecture to achieve this goal. The graphical illustration is shown in Figure 1.

The hybrid CNN-RNN architecture has been successfully applied to many applications including image segmentation Wang, Yang, Mao, Huang and Xu (2016a), speech emotion recognition Yao, Wang, Liu, Liu and Pan (2020). It provides a very natural way for feature extraction and statistical dependency modeling. As one part of CNN-RNN, CNN Lecun et al. (2015) is a specialized feedforward neural network, which is characterized by the presence of convolutional layers that use a stack of convolutional kernels to detect local patterns. Typically, a CNN consists of an input layer, multiple pairs of convolutional-pooling layers, a flatten layer, one or more fully connected layers, and the last softmax layer. The convolutional layer is the most crucial part of CNN. The output of a layer comes from its previous layer convolved with a set of filters, that is

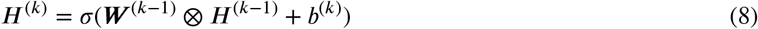

where *H*^(*k*)^, *W*^(*k*)^, and *b*^(*k*)^ respectively denote the feature map, convolutional filter, and biases of *k*-th layer, *H*^(0)^ = *E*_1_, *σ* denotes an activation function that is usually employed to guarantee the non-linearity of neural network. The most popular activation function is the rectified linear unit (ReLU) defined as *ReLU* (*x*) = max(0, *x*). In contrast, as shown in Figure 1, we employ a CNN architecture that receives two inputs, which separates two kinds of features (e.g., C2 encoding and kmer) by feeding them into additional univariate networks summed at the flatten layer of CNN, and then for a fixed window ***s***_*i,r*_, the flatten layer of CNN can be formulated as:

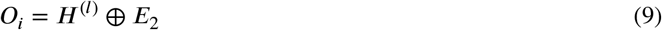

where *⊕* is the concatenation operator. *l* denotes the flatten layer of CNN. We adopt CNN to incorporate global sequence information and non-overlapping kmer features in viewing of its capabilities of modeling non-linearities and capturing local patterns such as codon. It is worth noting that this network architecture is very common in recent works Ghafoorian (2017); Zhehuan, Zhihao, Ling, Hongfei and Jian (2016) which claim that a better performance can be obtained when domain knowledge is incorporated into CNN.

The other part of CNN-RNN is an RNN which has been successfully applied to bioinformatics. Sequence data usually exhibit statistical dependencies and consider these dependencies usually yield performance benefits. There exists a considerable number of works that exert efforts to exploit statistical dependencies in DNA sequences, such as quantifying the function of DNA sequences Daniel and Xie (2016), subcellular protein localization Snderby, Snderby, Nielsen and Winther (2015), protein secondary structure prediction Spencer, Eickholt and Cheng (2015), segmentation of DNA sequences Cheng, Huang and Liou (2012). In our previous work Wei et al. (2020), we demonstrate that label dependencies among coding labels play an important role in protein coding regions prediction for transcript sequences. In this work, we extend this characteristic to genomic sequences and adopt the same BRNN architecture in Wei et al. (2020). Instead of estimating the conditional probability *p*(*y*_*i*_|***s***_*i,r*_, ***y***_¬*i,u*_) that has high-order dependencies among coding labels, the BRNN architecture in Wei et al. (2020) consider two-order dependencies among coding labels, and hence more computationally efficient, it reduces the problem of Eq. 4 to the following formula:

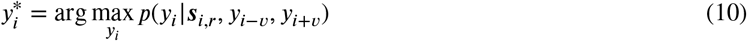

where *v* is a step interval that defines how far that two positions correlate. As shown in Figure 1, after obtaining the output of CNN for three subsequences, the forward and backward pass of BRNN can be formulated as:

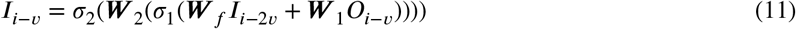

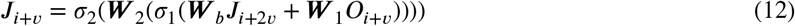

where ***W***_1_, ***W***_2_, ***W***_*b*_, ***W***_*f*_ respectively denote the weight matrices in the first hidden layer, second hidden layer, forward recurrent layer, backward recurrent layer of BRNN. *σ*_1_ and *σ*_2_ denote sigmoid and softmax activation function, respectively. ***I***_*i*_ and ***J***_*i*_ respectively denote the forward and backward passing message in a position *i* of a sequence. ***I***_0_ and ***J***_0_ denote the initial states with constant zero entries. From Eq. 11, we can see that the forward passing message ***I***_*i*_ in a position *i* is composed of the forward passing message in a position *i* − *v*, and the output of CNN in the position *i*. Similarly, the backward passing message ***J***_*i*_ in the position *i* is composed of the backward passing message in a position *i* + *v*, and the output of CNN in the position *i*. Actually, ***I***_*i*_ and ***J***_*i*_ is the estimations of *y*_*i*_, and the difference lies in that ***I***_*i*_ determines *y*_*i*_ by the current and past information whereas ***J***_*i*_ determines *y*_*i*_ by the current and future information. Finally, the prediction for the sample ***s***_*i,r*_ can be formulated as:

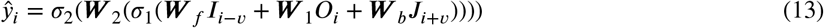

where 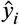 indicates how likely is it that the nucleotide in the center of the sliding window is coding. From Eq. 13, we can see that the prediction of a sample ***s***_*i,r*_ is dependent on the feedbacks from its neighboring positions *i* − *v* and *i* + *v*, and the output *O*_*i*_ of CNN in the position *i*.

Note that the step interval *v* must be a multiple of three for the reason that in practice the coding label sequence ***y*** is actually not always 1 in open reading frame, but shows a periodicity of three nucleotides (e.g., [1,0,0,1,0,0,…]). Meanwhile, as confirmed by experiments in Wei et al. (2020), the setting of *v* is significantly relevant to the prediction performance. Theoretically, the most neighboring positions (*v*=3) contribute the most to position *i*, while the reverse is true in our situation. If *v* is set small, ***s***_*i*_ is almost the same as ***s***_*i−v*_ and ***s***_*i*+*v*_, in which case ***I***_*i*−*v*_ and ***J***_*i*+*v*_ provide almost no additional information for the prediction 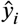. Hence, the setting of *v* is optimal when it equals the sliding window size, in which case ***s***_*i*_, ***s***_*i*−*v*_ and ***s***_*i*+*v*_ is completely different so that ***I***_*i*−*v*_ and ***J***_*i*+*v*_ provide the most information for the prediction 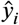.

## 4. Result

In this section, we conduct three experiments on four gene datasets. The first is to prove the significance of hybrid encoding. In the second experiment, we make a comparison of the proposed method to existing state-of-art methods such as C4+MLP Hatzigeorgiou et al. (1996), C4+SVM Shuo and Yi-sheng (2009), Z curve+LDA Zhang et al. (1998), kmer+MLP Guigó (1997); Hatzigeorgiou (2002), kmer+SVM Tzanis et al. (2012), kmer+BRNN Wei et al. (2020). The goal of the last experiment is to evaluate the time cost of the proposed method. The network architectures are shown in Table 1.

**Table 1.**
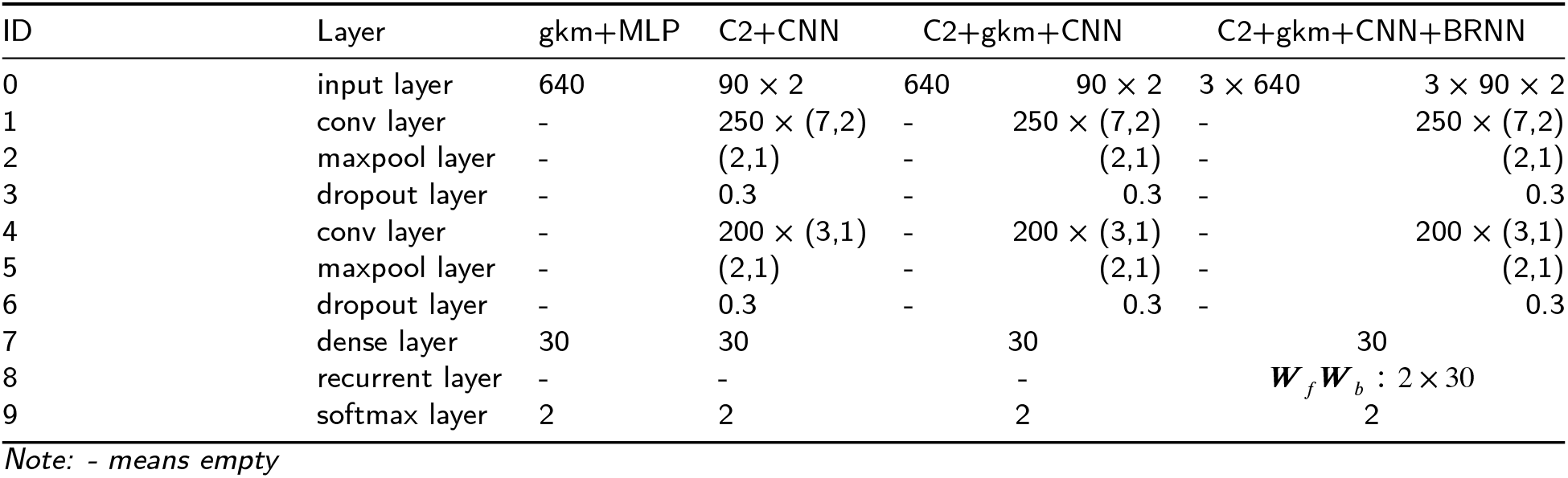
Different network architectures in experiments.

### 4.1. Performance measurements

In order to evaluate the performance of the proposed method for coding regions prediction, the analysis in this paper employs three evaluation criteria in terms of Sensitivity (Sn), Specificities (Sp) and Area Under the Receiver Operating Characteristic curve (auROC). All these criteria are based on the notions of TP, FP, TN, and FN, which correspond to number of true positives, false positives, true negatives, and false negatives. In a ROC, one typically plots the true positive rate (TPR=TP/(TP+FN)) as a function of the false negative rate (FNR=FN/(FN+TN)). The auROC can be calculated by using the trapezoidal areas created between each ROC points. The detailed definition can be found in Mitchell, Carbonell and Michalski (1997); Davis and Goadrich (2006).

### 4.2. Significance of hybird encoding

In order to prove the effectiveness of hybrid encoding, we conduct an ablation study to separate the hybrid encoding and observe the prediction performance for each single encoding. To be specific, hybrid encoding is separated into C2 and gkm encoding and then fed into CNN and MLP (e.g., C2+CNN and gkm+MLP). As it can be seen from Table 2-3, C2+gkm+CNN significantly outperform C2+CNN and kmer+MLP, both on genomic and transcript sequences. Moreover, it is worth noting that C2 encoding achieves better performance than kmer+MLP and gkm+MLP in genomic sequences, which prove the effectiveness of global sequence order information, but at the cost of computational complexity. Moreover, gkm+MLP achieves a little better performance than C2+CNN in transcript sequences. Also it achieves much better performance than kmer+MLP, which proves that larger *k* can provide more useful information than using *k* = 3. All the prediction performance of the above methods decreases from transcript sequences to genomic sequences for the fact that coding regions are continuous on transcript sequences but interrupted by introns on genomic sequences. In brief, we can conclude from the result that there exists a complementation relationship between C2 encoding and gkm features, integration of which can facilitate the machine learning method to fully capture coding features.

**Table 2.** Performance comparison of our proposed method with the other state-of-the-art methods on genomic datasets.

| Method | Human |  |  | Mouse |  |  |
| --- | --- | --- | --- | --- | --- | --- |
|  | Sn | Sp | auROC | Sn | Sp | auROC |
| Z curve+LDA Zhang et al. (1998) | .8010 | .8462 | - | .7829 | .8114 | - |
| C4+MLP Hatzigeorgiou et al. (1996) | .8993 | .8900 | .9615 | .9032 | .8708 | .9558 |
| C4+SVM Shuo and Yi-sheng (2009) | .8609 | .8629 | - | .8369 | .8369 | - |
| kmer+MLP Guigó (1997) | .8528 | .8836 | .9413 | .8645 | .8723 | .9425 |
| gkm+MLP | .8831 | .8977 | .9588 | .8775 | .8957 | .9562 |
| C2+CNN | .9406 | .9043 | .9772 | .9445 | .8922 | .9755 |
| C2+gkm+CNN | .9227 | .9360 | .9808 | .9256 | .9260 | .9789 |
| C2+gkm+CNN+BRNN | **.9621** | **.9631** | **.9930** | **.9635** | **.9598** | **.9926** |

**Table 3.** Performance comparison of our proposed method with the other state-of-the-art methods on transcript datasets.

| Method | Human |  |  | Mouse |  |  |
| --- | --- | --- | --- | --- | --- | --- |
|  | Sn | Sp | auROC | Sn | Sp | auROC |
| Z curve+LDA Zhang et al. (1998) | .8131 | .8407 | - | .8339 | .8422 | - |
| kmer+MLP Hatzigeorgiou (2002) | .9479 | .9172 | .9810 | .9403 | .9180 | .9793 |
| kmer+SVM Tzanis et al. (2012) | .9276 | .9292 | - | .9291 | .9271 | - |
| kmer+BRNN Wei et al. (2020) | .9825 | .9739 | .9975 | .9793 | .9791 | .9973 |
| gkm+MLP | .9416 | .9462 | .9864 | .9571 | .9237 | .9851 |
| C2+CNN | .9491 | .9323 | .9856 | .9329 | .9352 | .9826 |
| C2+gkm+CNN | .9595 | .9360 | .9885 | .9563 | .9330 | .9870 |
| C2+gkm+CNN+BRNN | **.9908** | **.9797** | **.9986** | **.9910** | **.9814** | **.9985** |

**Table 4.** Brief description of time cost on four datasets with regard to the proposed method.

| Dataset | Type | Number of sequences | Number of codons | Time Cost (s/epoch) |
| --- | --- | --- | --- | --- |
| Human | genomic | 19,288 | 6,809,025 | 4,340 |
| Mouse | genomic | 16,473 | 5,787,480 | 3,684 |
| Human | transcript | 24,842 | 7,855,557 | 5,072 |
| Mouse | transcript | 19,900 | 6,549,798 | 4,104 |

### 4.3. Performance comparison with existing state-of-the art methods on genomic and transcript sequences

We compare the performance of the proposed method with that of existing methods such as C4+MLP, C4+SVM, Z curve+LDA, kmer+MLP, kmer+SVM, kmer+BRNN. All the methods are trained and evaluated with the same dataset for a fair comparison. From Table 1-2, it is observed that our proposed method performs the best among the existing methods and achieves the highest Sn, Sp and auROC scores on all the four datasets, the average Sn, Sp, auROC scores reach .9628, .9615, .9928 on genomic datasets, and .9909, .9806, .9985 on transcript datasets, respectively. We also plot the ROC curve on genomic and transcript sequences. As it can be seen in Figure 2, given a fixed false positive rate of .05, our proposed method respectively achieves an average sensitivity of .9720 on genomic datasets, an improvement of .2021 over the second best method, C4+MLP. Meanwhile, given a fixed false positive rate of .01, our proposed method respectively achieves an average sensitivity of .9805 on transcript datasets, an improvement of .0224 over the second best method, kmer+BRNN. All the results demonstrate that our proposed method is a high-accuracy protein coding regions prediction method, especially in transcript sequences.

**Figure 2:**
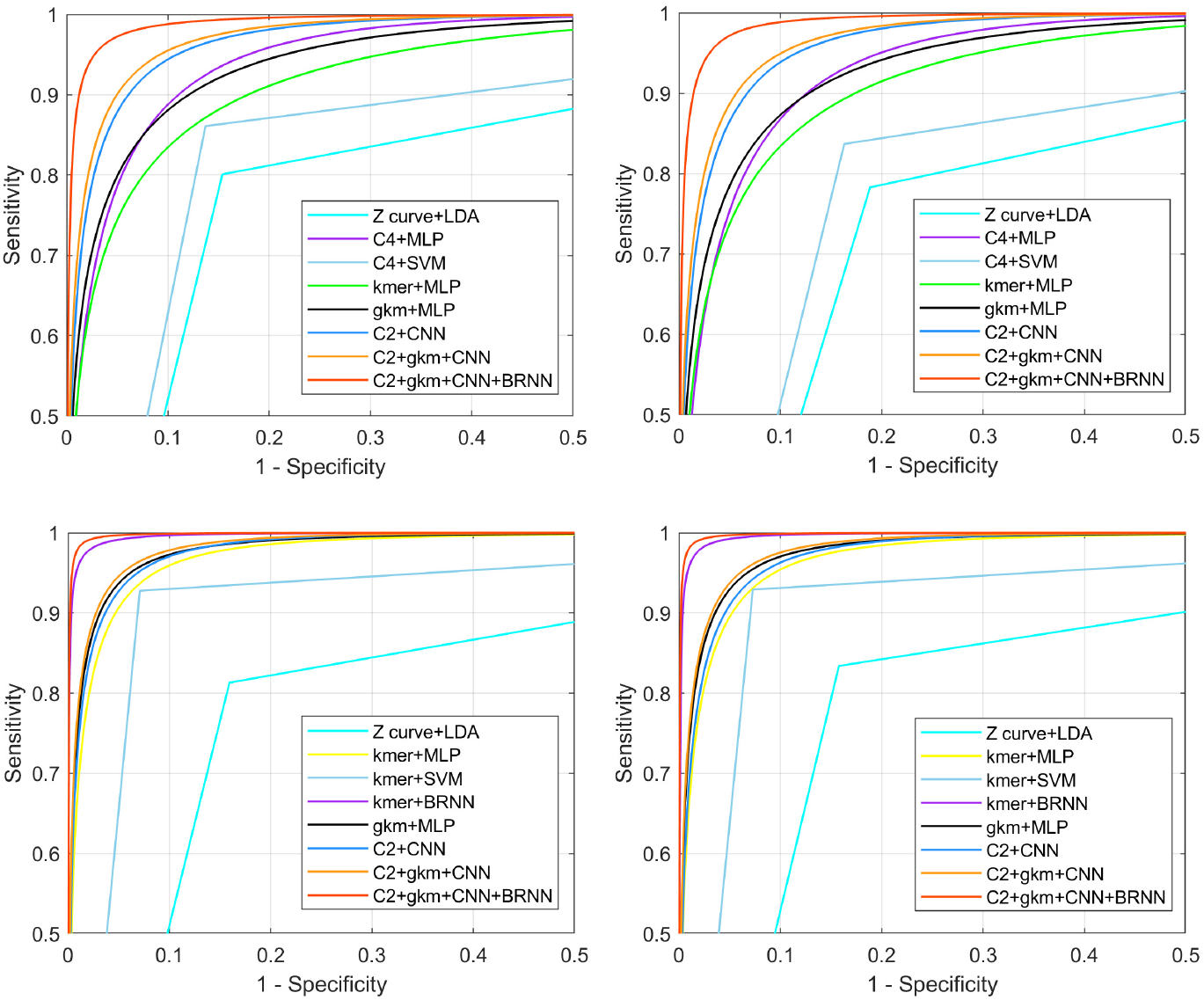
Performance comparison of the proposed method with C4+MLP, C4+SVM, Z curve+LDA, kmer+MLP, kmer+SVM and kmer+BRNN. (***A***) the ROC curves on genomic Human dataset; (***B***) the ROC curves on genomic Mouse dataset; (***C***) the ROC curves on transcript Human dataset; (***D***) the ROC curves on transcript Mouse dataset.

### 4.4. Time cost of the proposed method

We further briefly analyze the computational cost of the proposed method. All the experiments are conducted on an Intel Core i5-10400 CPU 2.90 GHz PC with 16 GB RAM. The proposed method is implemented mainly in Tensorflow and partly in Matlab. Table 4 gives the time cost of the proposed method on four datasets.

## 5. Conclusions

In summary, protein coding regions prediction is a very important but overlooked subtask for tasks such as prediction of complete gene structure, coding/non-coding RNA. However, it is still a lack of effective computational methods to learn coding features from genomic and transcript sequences. Indeed, coding features in biological sequences usually exhibit heterogeneity (e.g., global sequence order information, frequency domain of features like kmer, statistical dependencies among coding labels) and are difficult to capture by using a single encoding scheme and machine learning method. In this paper, we present a deep learning framework with hybrid encoding for protein coding regions prediction, which effectively incorporates the three kinds of features into a hybrid CNN-BRNN architecture and achieves a remarkable prediction performance when compared with existing state-of-the-art methods on genomic and transcript sequences.

## Acknowledgments

This work is supported by the Natural Science Foundation of China under Grants 11674352, and 91853123.

